# Hemoperfusion of pigs with a carbon-cellulose cartridge: a pilot study revealing a new animal model of extended anaphylactic shock

**DOI:** 10.64898/2026.03.05.709860

**Authors:** Bálint A. Barta, Tamás Radovits, Sylvia Spiesshofer, Albert J. Husznai, Attila B. Dobos, László Mészáros, Tamás Mészáros, Gergely T. Kozma, Réka Facskó, László Dézsi, János Nacsa, Joshua A. Jackman, Béla Merkely, János Szebeni

## Abstract

To investigate the immune mechanisms underlying extracorporeal circulation-associated anaphylactoid reactions, we inserted externally perfused cartridges into the venous circulation of pigs, including a cellulose-coated activated-charcoal adsorbent (Adsorba^®^ 300C), a polysulfone hollow-fiber membrane hemofilter, and polypropylene hollow-fiber-based heart-lung machine gas-exchange oxygenators. Blood was circulated using a roller pump, and the animals were monitored for systemic and pulmonary arterial pressures (SAP, PAP) changes, blood levels of complement C3a, thromboxane B_2_, and hemoglobin, blood cell counts and hematocrit. None of the cartridges caused major changes in these endpoints except the Adsorba^®^ 300C, which displayed a fulminant anaphylactoid reaction characterized by profound hypotension, maximal pulmonary hypertension, hemoconcentration, thrombocytopenia and a surge of C3a and thromboxane B_2_, i.e., hallmarks of complement activation-related pseudoallergy. Within 15 min, the reaction advanced to profound hemodynamic collapse, which was managed with norepinephrine and cardiopulmonary resuscitation. After brief rebound hypertension, shock recurred despite repeated rounds of resuscitation until death. Considering that major adverse reactions with overlapping symptoms have been reported in humans subjected to hemoperfusion with the same cartridge, this porcine model provides clinically relevant insights into the mechanisms of extracorporeal circulation-induced anaphylactoid responses. Furthermore, the Adsorba^®^ 300C-induced physiological changes along the immune-cardiopulmonary axis represents a new animal model for irreversible cardiovascular collapse escalating into shock.

## 1. Introduction

Carbon-cellulose hemoperfusion cartridges, such as Adsorba^®^ 300C, are effective toxin-removal devices [1, 2] associated with several well-recognized adverse events (AEs) [3-6]. The most frequent and consistent pathological changes include hypotension, tachycardia, transient leukopenia, rapid thrombocytopenia, and a bleeding tendency, particularly in allergy-prone susceptible patients. Collectively, these phenomena reflect the comparatively lower biocompatibility of older carbon-based sorbents relative to modern polymeric hemoperfusion cartridges. Table 1 categorizes the symptoms by organ system and highlights the mechanisms contributing to their development.

**Table 1.**
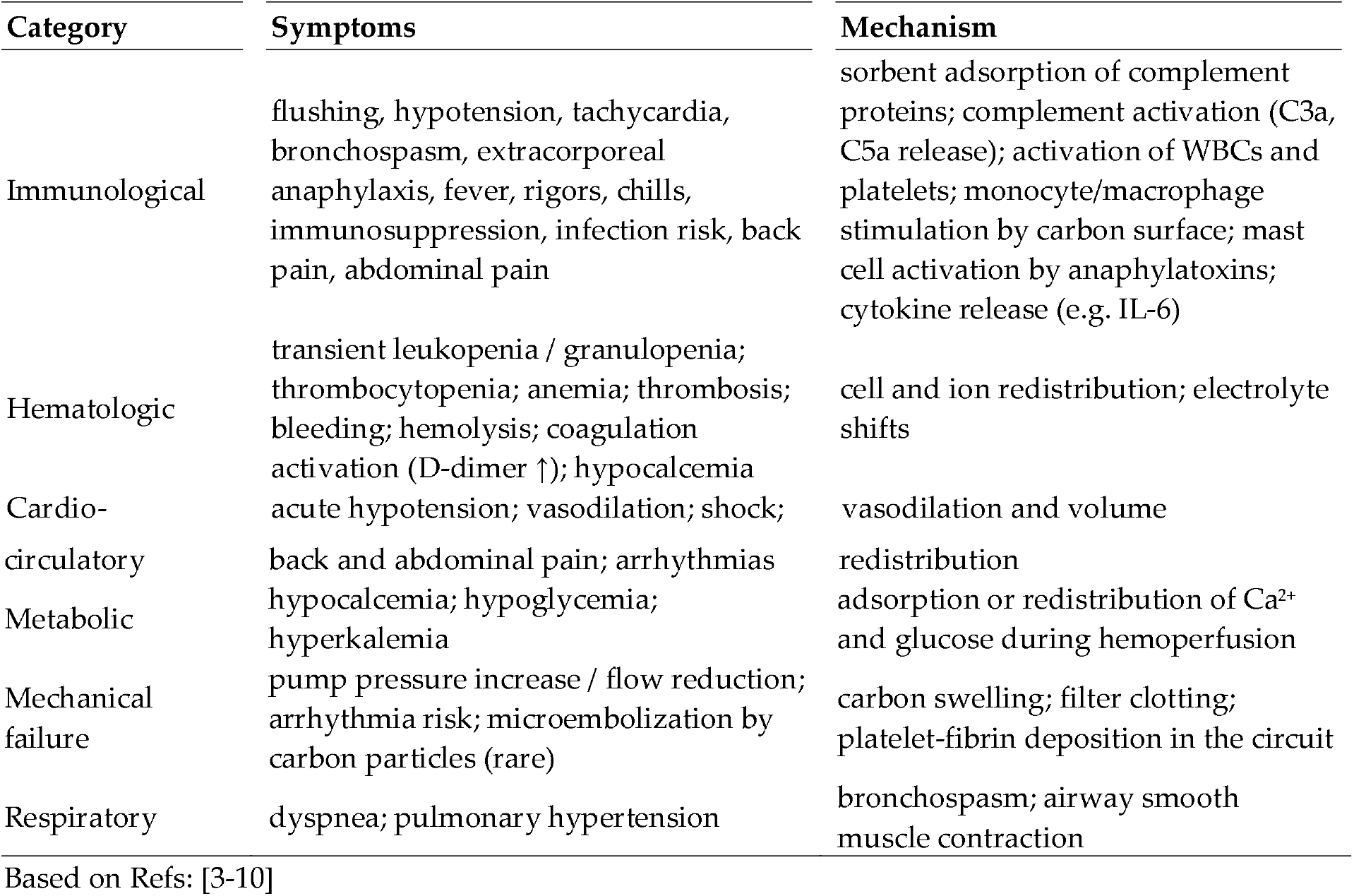
Adverse events caused by hemoperfusion in humans.

Although generally manageable, these AEs require strict monitoring of blood pressure, platelet count, metabolic parameters, and other physiological markers during treatment, which is a major hurdle in their clinical use. Thus, strategies to mitigate or prevent these AEs represent an unmet medical need, one that necessitates the development and use of an appropriate animal model.

As shown in Table 1, activation of the complement (C) system, the humoral arm of innate immunity, is a pivotal immune mechanism of perfusion reactions. The phenomenon corresponds to C activation-related pseudoallergy (CARPA), an acute hypersensitivity reaction that typically arises following the intravenous (i.v.) administration of certain nanomedicines or other complex drugs [11,12]. A well-established preclinical model for studying CARPA is the porcine model, as pigs exhibit high and quantitatively reproducible cardiopulmonary sensitivity to complement-mediated reactions [13–18].

The aim of this study was to explore the use of this model to investigate extracorporeal adsorber– induced cardiopulmonary stress reactions along the immune–circulatory system axis, i.e., the core pathophysiological features of CARPA. In pigs, these reactions are characterized by pulmonary hypertension, with or without systemic hypotension, leukopenia, granulocytopenia, thrombocytopenia, hemoconcentration, and skin flushing or rash, closely mimicking severe anaphylactoid reactions in humans [13, 16-18]. An additional advantage of the porcine model is that it accommodates full-size clinical devices, making it suitable for predicting human reactogenicity and evaluating mitigation or therapeutic approaches.

## 2. Materials and Methods

### 2.1. Extracorporeal cartridges

The extracorporeal hemoperfusion adsorber cartridge (Adsorba® 300C) and the hollow-fiber membrane hemofilter (multiFiltratePRO) were manufactured by Fresenius Medical Care GmbH (Hamburg, Germany), whereas the heart–lung machine oxygenator (Inspire 8 Start P) was manufactured by Sorin Group Italia S.r.l. (Mirandola, Italy). The carbon (activated charcoal) particles in Adsorba^®^ 300C are coated with cellulose acetate polymer, which reduces platelet adhesion, hemolysis, and excessive protein loss while maintaining high adsorption capacity for small-and medium-molecular-weight toxins [19]. The hemofilter (multiFiltratePRO) were manufactured by Fresenius Medical Care GmbH (Hamburg, Germany), and the heart-lung machine oxygenator (Inspire 8 Start P) was from by Sorin Group Italia S.r.l. (Mirandola, Italy). The functions, composition, adsorption/membrane surface areas and other features of all these cartridges are detailed in Table 3 in the text.

### 2.2. Animals

Pig experiments were performed under ethical approvals PE/EA/00246-2/2023 and PE/EA/00531-6/2025. SPF Danbred pigs of both sexes (2-3 months old, 20-28 kg) were obtained Sano-Pikk Ltd., (Zsámbék, Hungary). The pigs were housed in a vivarium at a temperature of 22 °C, a humidity of 33%, and a 12-hour light/dark cycle. The animals were allowed to drink ad libitum and were fed 400 grams of preformulated pig feed twice daily. The animals were subjected to a 12-hour fasting period before the procedure.

Before surgery, the pigs were premedicated with an intramuscular (IM) injection of 0.25 mg/kg midazolam, 0.2 mg/kg xylazine, and 10 mg/kg ketamine. Peripheral vascular access was obtained via the left lateral auricular vein. Anesthesia was induced using a face mask delivering isoflurane 5 vol% in 100% FiO_2_. After orotracheal intubation, anesthesia was maintained with 2 vol% isoflurane in 100% FiO_2_.

Pigs were cannulated in the right femoral vein for blood sampling using a 6 Fr sheath, and in the left femoral artery for mean arterial pressure (MAP) measurement using a DOME-type invasive blood pressure transducer (6 Fr). The right internal jugular vein was cannulated for insertion of a Swan-Ganz catheter to measure pulmonary arterial pressure (PAP) (6 Fr). The left jugular vein was connected to the inlet of the extracorporeal hemoperfusion apparatus (ECHA; Watson-Marlow Ltd., UK), consisting of a peristaltic roller pump that provided continuous, non-pulsatile blood flow at adjustable speed control (10-200 rpm) and medical-grade silicone tubing for extracorporeal circulation. The outlet of the pump was connected to a hemoperfusion cartridge (Adsorba^®^ 300C) using dialysis catheter tubes (JOLINE, 13.5 Fr, 150 mm; ref. PKTHF13P150; LOT 502033). The cartridge was flushed with 2 L of heparinized saline before starting the experiment.

Whole blood is circulated extracorporeally through the cellulose-coated charcoal bead-filled cartridge using a motor-driven roller pump. During the experiment, standard CARPA monitoring parameters were recorded, including systemic and pulmonary arterial pressure (SAP, PAP), blood cell counts. Mean SAP was measured in the left femoral artery via a DOME-type invasive transducer (6 Fr). After obtaining the baseline (negative control, 5 mL Salsol infusion), hemoperfusion was initiated and the flow was increased gradually until reaching 0.15 L/min. at which rate blood passed through the cartridge and returned to the animal within approximately 3.5 minutes. At the end of hemoperfusion, blood remaining in the cartridge was washed back into the animal’s circulation using heparinized Salsol at 0.20-0.25 L/min, until the effluent fluid became clear.

Figure 1 presents a schematic overview of the porcine instrumentation and extracorporeal circuit used for hemoperfusion along with light micrographs of the cartridge’s inner section and the charcoal-beads.

**Figure 1.**
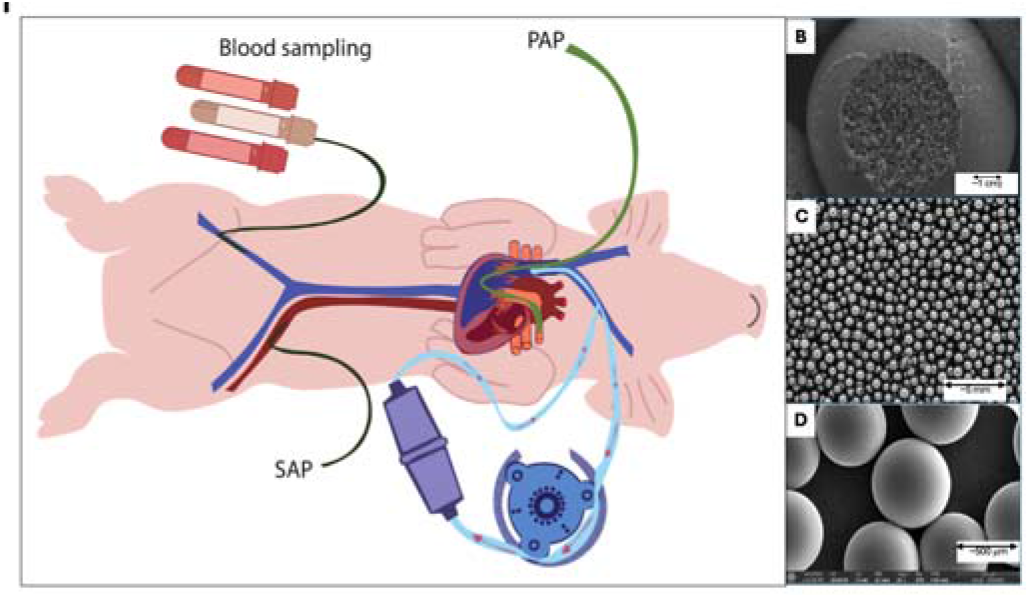
A) Layout of the pig experiment using Adsorba^®^ 300C and other cartridges. Simultaneous monitoring of systemic arterial pressure (SAP) and pulmonary arterial pressure (PAP) were performed invasively by arterial cannulation and Swan-Ganz catheterization, respectively. B Cross section of the Adsorba^®^ 300C cartridge. C, D, Close-up images of the microbeads within the column (B-D). Images obtained from publicly accessible online sources and included for illustrative purposes only. Bead diameter: 0.3-0.8 mm. Unlike hemodialysis, hemoperfusion relies on surface adsorption rather than diffusion across a membrane [20]. The column binds circulating toxins, drugs, complement fragments, cytokines, or other inflammatory mediators by direct adsorption onto the huge sorbent surface via hydrophobic, ionic, or affinity interactions. The purified blood is then returned to the circulation.

### 2.4. Blood sampling and hematology

Peripheral blood was collected every 10 min during hemoperfusion, via indwelling vascular catheters into K_2_EDTA-anticoagulated tubes and analyzed within 1 h. Hematological measurements were performed using an automated veterinary hematology analyzer (Abacus®, Diatron, Budapest, Hungary) operated with porcine-appropriate settings and calibrated according to the manufacturer’s instructions.

### 2.5. Blood cell analysis

Blood cell counting was based on electrical impedance (Coulter principle) and photometric hemoglobin determination. The automated standard blood panel provided white blood cell (WBC), red blood cell (RBC), platelet (PLT) counts, hemoglobin (Hb), hematocrit (Ht), mean corpuscular volume (MCV), mean corpuscular hemoglobin (MCH), and mean corpuscular hemoglobin concentration (MCHC). Cellular elements were quantified by impedance counting, Hb was measured photometrically after erythrocyte lysis, and hematocrit was calculated from RBC count and MCV.

### 2.6. Measurement of C3a and Thromboxane B2 (TXB2)

Blood samples were collected every 10 min into EDTA tubes kept on ice, followed by centrifugation at 3,000 × g for 10 min at 4 °C. Plasma aliquots were stored at-70 °C until analysis. The C activation product sC5b-9 and thromboxane B_2_ (TXB_2_) in plasma were measured by enzyme-linked immunosorbent assay (ELISA), as described previously [17]. Blood samples for TXB_2_ analysis were also supplemented with indomethacin (5 μg/mL) to inhibit ex vivo cyclooxygenase activity. Absorbance in ELISA wells was measured using a microplate reader, and analyte concentrations were calculated from standard curves.

## 3. Results

### 3.1. Hemodynamic changes during extracorporeal circulation of PBS as negative control

To provide background reference for the effects of Adsorba^®^ 300C and other cartridges tested in our model we have explored the hemodynamic effects of motor-assisted extracorporeal circulation per se, without any connected cartridge. Figure 2 shows the changes of systemic arterial pressure (SAP, red) and pulmonary arterial pressure (PAP, blue) during a 60 min of motor-assisted circulation and rinsing in the tube content into the circulation.

**Figure 2.**
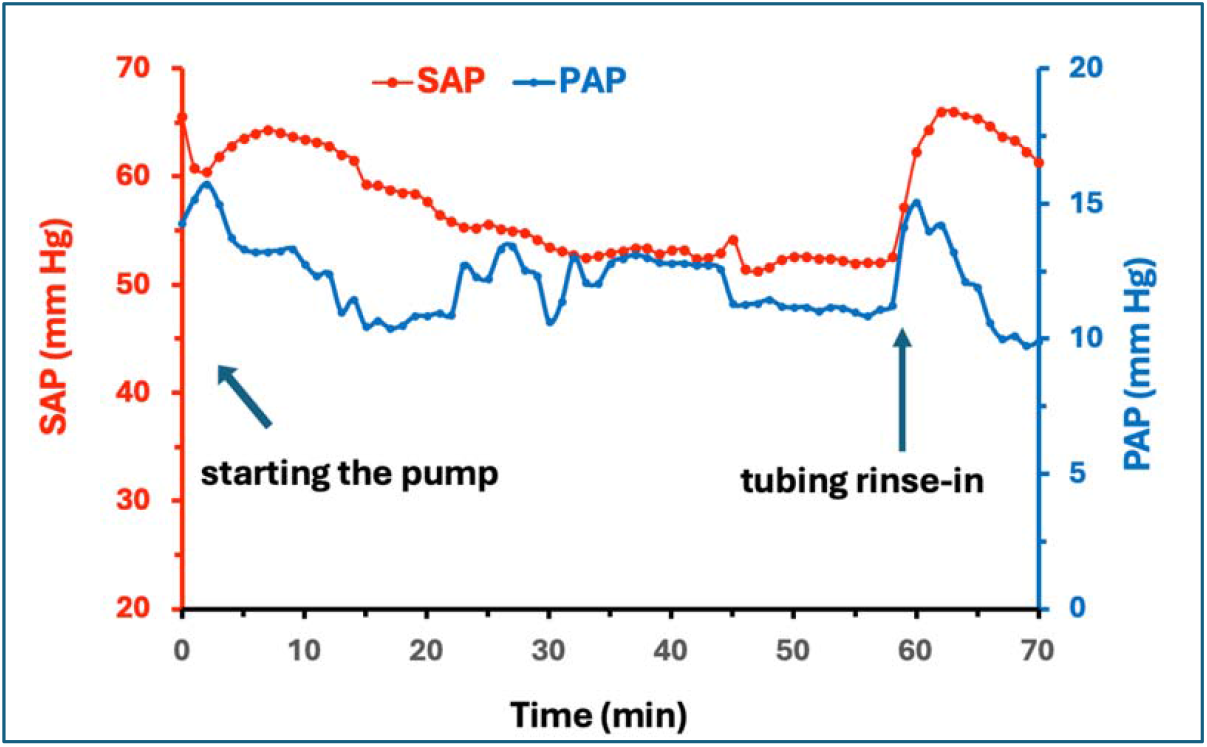
Hemodynamic effects of roller pump-driven extracorporeal circulation. A pig was instrumented identically to hemoperfusion experiments, except that the adsorbent cartridge was omitted from the extracorporeal loop. Continuous recordings with minute-level readouts.

The curves revealed an initial drop, followed by a gradually developing, approximately 20 mmHg overall drop of SAP during the pump operation. Normal blood pressure was then restored by rinsing the tube content back into the circulation. Although it has not been investigated, the pumping-caused hypotension could be a combination of extra blood volume due to the tubing and overall changes in the blood flow, possibly influencing the baroreceptors in the carotid sinus and the aortic arch. The restoration of SAP at the end of the experiment is consistent with the impact of blood volume. In contrast, the changes of PAP between 10-15 mm Hg were negligible during pump operation. Overall, the above pump-induced hemodynamic changes were minor and limited.

### 3.2. Systemic and pulmonary pressure changes during Adsorba^®^ 300C hemoperfusion

Figure 3 details the dynamics of blood pressure changes following initiation of extracorporeal circulation, first for 20 min with PBS control, and subsequently after insertion of the Adsorba® 300C charcoal matrix into the circuit (approximately after baseline). PBS caused no significant changes in blood pressure (Fig. 3A,B), whereas circulation through the charcoal cartridge induced marked hemodynamic alterations. After ∼10 min of perfusion, an abrupt decline in SAP occurred, progressing to overt shock within approximately 8 min (Fig. 3C). At this point, a combination of intravenous adrenaline (EPI) and norepinephrine (NE), together with cardiopulmonary resuscitation (CPR) had to be applied to salvage the animal. This intervention transiently elevated SAP to ∼170% of baseline within 10 min, indicating a compensatory blood pressure overshoot. However, unlike in many previous porcine studies in which the hemodynamic effects of various nanoparticles, diagnostic agents, and complex macromolecules were tested, any observed decline in SAP could be reliably normalized after reanimation [18]. In the present case, however, despite repeated resuscitation, the blood pressure continued to fall below the shock threshold, indicating sustained progression toward unavoidable shock and death.

**Figure 3.**
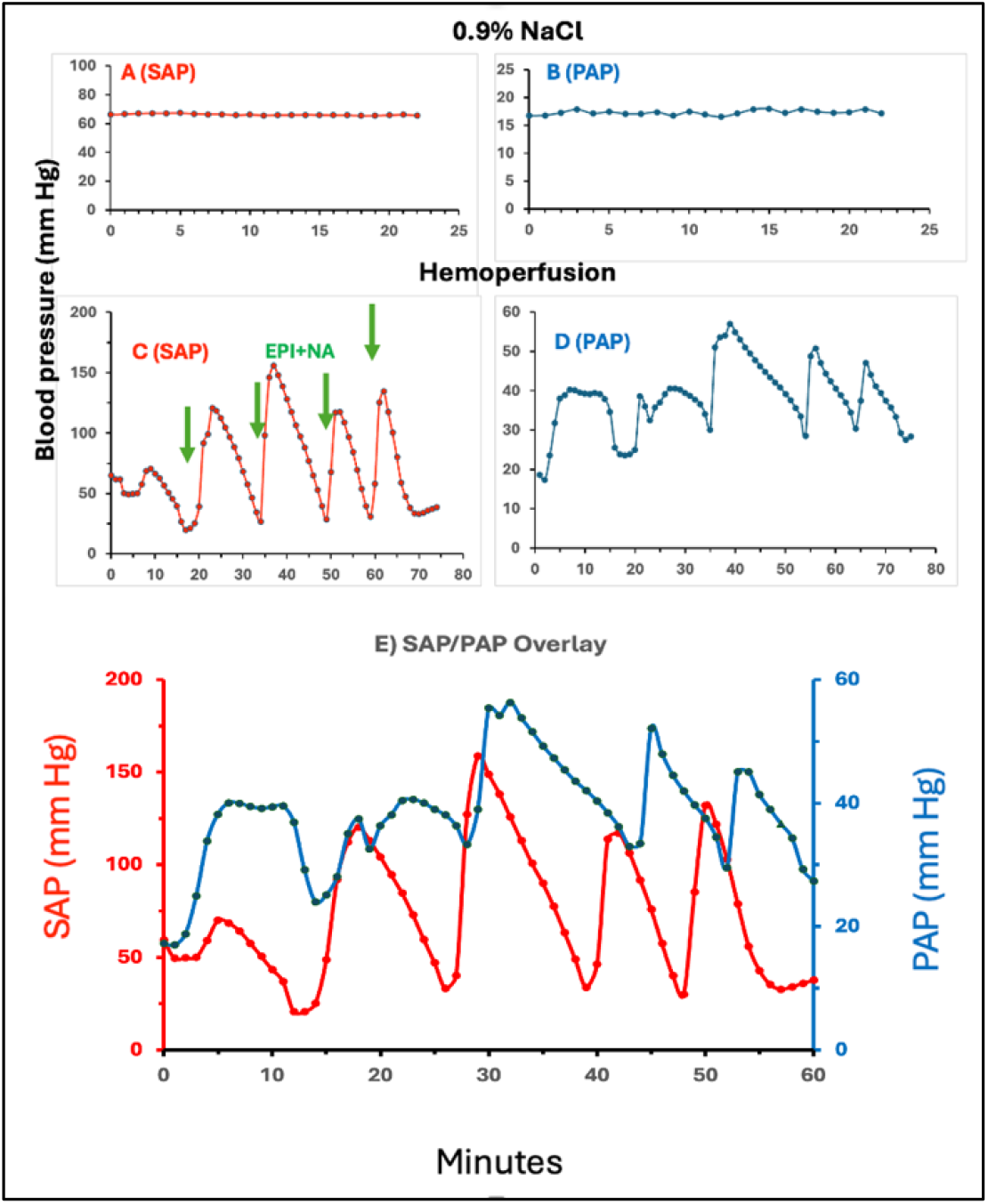
Systemic and pulmonary blood pressure changes in a pig during hemoperfusion using the Adsorba 300C cartridge. Mean arterial pressure (MAP) was measured in the left femoral artery using an invasive blood pressure transducer, while pulmonary arterial pressure (PAP) was measured via a Swan-Ganz catheter inserted through the right internal jugular vein and positioned in the pulmonary artery.

The PAP exhibited comparable oscillatory peaks and troughs (Fig. 3D), which, as shown in Fig. 3E, were preceded by SAP peaks by several minutes after the second oscillation. This time lag between the SAP and PAP peaks is consistent with an anatomically based causal relationship wherein the rise in SAP led to subsequent increase in PAP due to enhanced filling of the pulmonary artery, whereas the rise in PAP has contributed to a decline in SAP by impairing left ventricular filling secondary to pulmonary vasoconstriction.

Taken together, the above experiment revealed a new large-animal model of the severe toxicity of the hemoperfusion column. Compared with the acute, fulminant anaphylactic shock triggered by bolus injection of nanoparticles [13, 14, 16-18].

### 3.3. Effects of oxygenator and dialysator cartridges on blood pressure of pigs

To compare the above effects with those induced by other types of extracorporeal circuits, we examined the impacts of additional ECHAs, including a hollow fiber membrane oxygenator (e.g., polyester microporous membrane oxygenator and a hemodialysis column (e.g., polysulfone-based high-flux dialyzer). Unlike the adsorption mechanism of Adsorba^®^ 300C and hemoperfusion cartridges, these devices operate via diffusion of gases (oxygenator) or small solutes (dialyzer) across semipermeable membranes [20]. The experiments were performed under conditions like those described above for the hemoperfusion studies. To assess inter-individual variability, 3 identical oxygenator columns were tested in three pigs. As shown in Fig. 4A and B, these columns also induced oscillations in PAP and SAP; however, these changes were markedly less pronounced. In the case of the oxygenators, the individual variation was minimal and remained within background noise levels.

**Figure 4.**
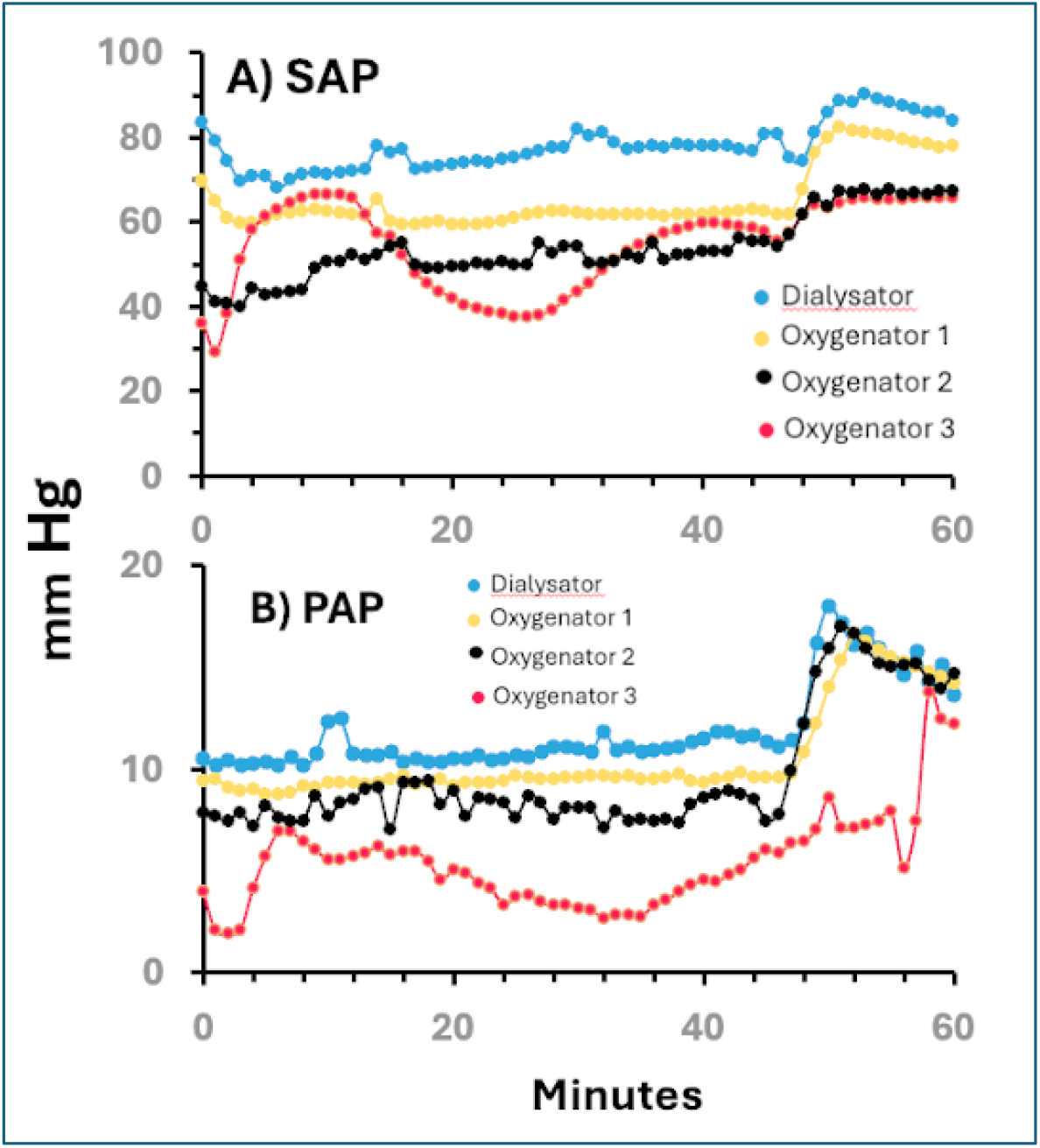
Systemic and pulmonary blood pressure changes in pigs during extracorporeal circulation using 3 heart-lung machine oxygenators (Oxygenator 1-3, Inspire 8 Start P) and 1 dialysis cartridge (multiFiltratePRO). Their sources are given in the Methods part. All experimental details are the same as in Fig. 3.

### 3.4. Hematological changes during extracorporeal circulation using different cartridges

#### 3.4.1. Extracorporeal circulation without cartridge

Figure 5 shows that the motor-aided extracorporeal blood circulation did not induce major hematologic alterations, apart from a moderate granulocytosis with concomitant trend for lymphopenia, possible consequences of mild immune activation. The apparent anemic trend with minor decrease of Ht is consistent with hemodilution by the extracorporeal circuit.

**Figure 5.**
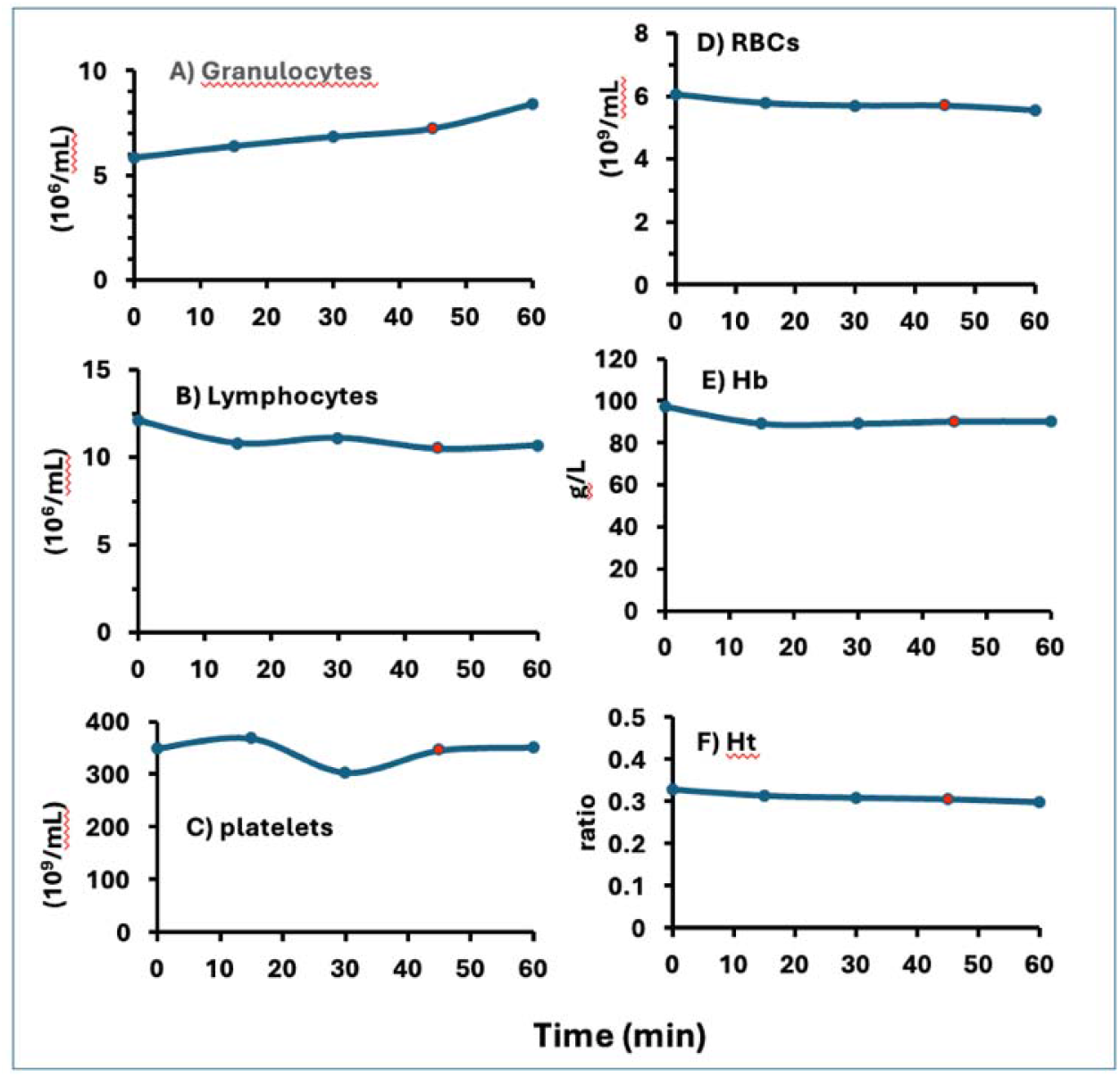
Effects of roller pump-driven extracorporeal circulation on hematology parameters. A pig was instrumented identically to hemoperfusion experiments, except that the adsorbent cartridge was omitted from the extracorporeal loop. Cell count, Hb and Ht determinations were done as described in the Methods.

#### 3.4.2. Adsorba^®^ 300C

Unlike the PBS control shown in Fig. 5, insertion of the Adsorba^®^ 300C cartridge into the extracorporeal circuit induced marked alterations in blood cell parameters, affecting both the white blood cell (WBC) and red blood cell (RBC) compartments. These changes included pronounced and persistent leukopenia (Fig. 6A), primarily attributable to granulopenia (Fig. 6B), transient lymphopenia (Fig. 6C), and mild thrombocytopenia (Fig. 6D). In parallel, there was a substantial increase in RBC count (Fig. 6E), hemoglobin concentration (Fig. 6F), and hematocrit (Fig. 6G). The WBC alterations are consistent with a fulminant inflammatory response, whereas the elevations in RBC-related parameters indicate hemoconcentration, likely reflecting increased capillary permeability and plasma extravasation.

**Figure 6.**
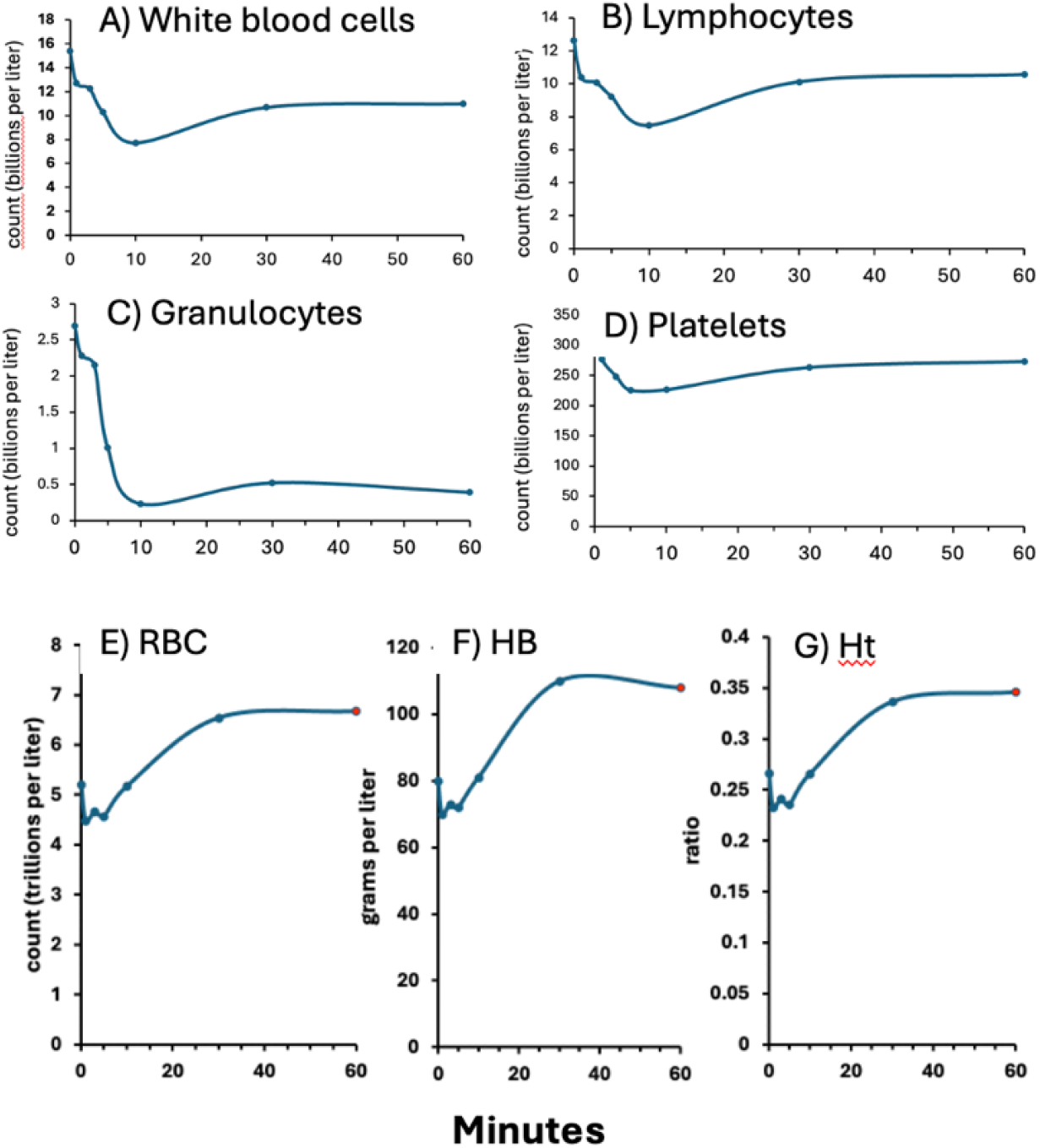
Effects of extracorporeal circulation with Adsorba 300C cartridge on hematology parameters. The same experiment as shown in Fig 3. Cell count, Hb and Ht determinations were done as described in the Methods. HB, hemoglobin, Ht, hematocrit

As for the volume of edema in the interstitial space and lung, a 28% increase in Ht while the RBC count is diluted in the EC tubing and the RBC mass remains unchanged (i.e., no bleeding), corresponds to a minimum of 22% reduction in total blood volume, entirely attributable to plasma loss. This implies, by rough estimation, a 31–37% reduction of the initial plasma volume, which in a 25-kg pig corresponds to approximately 300–370 mL of plasma. Such major imbalance in fluid homeostasis most likely represents a major contributor to shock and death.

### 3.5. Hemoperfusion using Adsorba^®^ 300C cartridge activates the humoral and cellular innate immune system

Figure 7 shows that perfusion of blood through the hemoperfusion cartridge induced a rapid and steep rise in C3a within the first minutes of circulation, indicating strong C activation. The early C3a peak then gradually declined despite ongoing perfusion, consistent with the rapid metabolism and cellular uptake of C3a. The parallel, nearly thousand-fold increase in TXB_2_ was also exceptionally pronounced, displaying delayed kinetics and reaching maximal levels later than C3a. This delay is consistent with secondary platelet and leukocyte activation downstream of C activation. The temporal dissociation between the C3a and TXB_2_ responses indicates a primary C-driven trigger of innate immune cell activation, reflecting sequential engagement of the humoral and cellular arms of the innate immune system.

**Figure 7.**
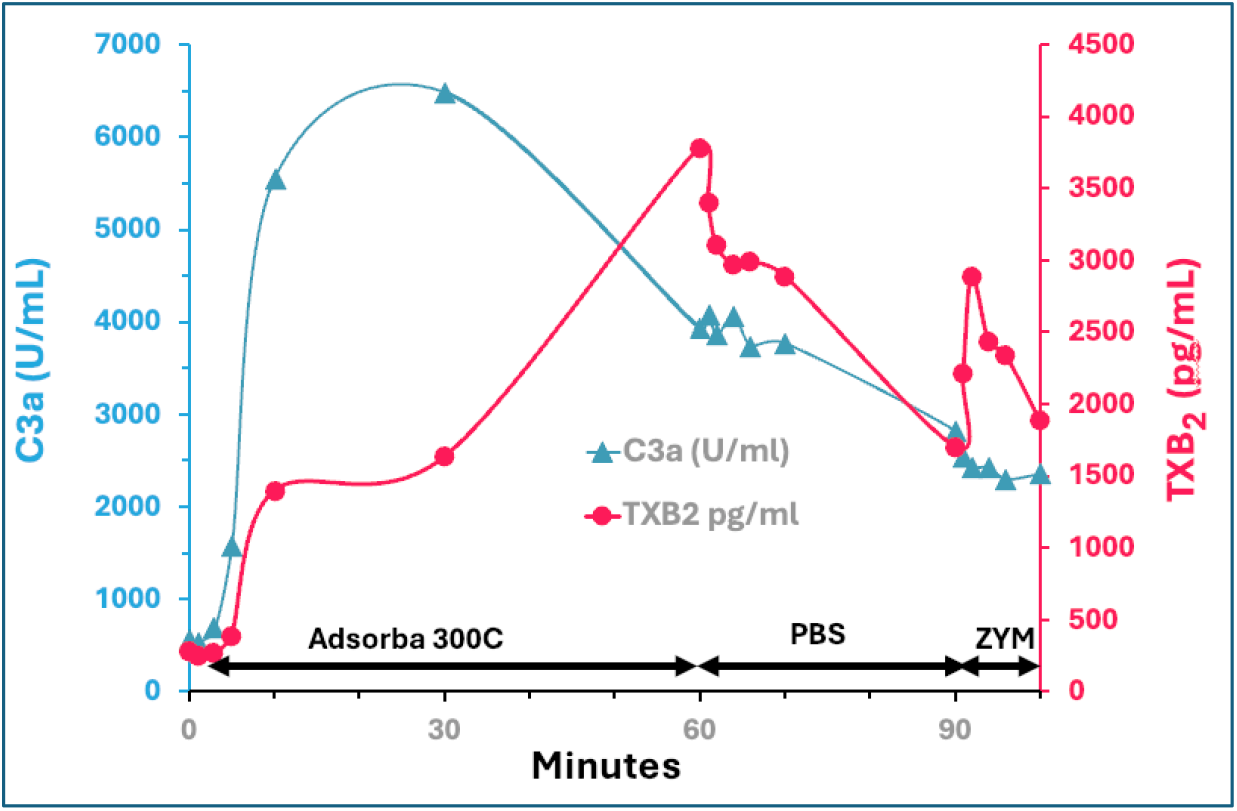
Complement activation and thromboxane release during hemoperfusion using an Adsorba^®^ 300C cartridge. Time course of plasma C3a (blue symbols, U/mL) and thromboxane B_2_ (TXB_2_; red symbols, pg/mL). Both parameters were measured in blood samples obtained at the indicated time points by ELISAs. Of note, the hemodynamic changes were monitored only for 60 min, during the hemoperfusion, while these samples were taken until the end of the experiment.

Upon initiation of extracorporeal circulation, plasma proteins rapidly adsorb to the activated carbon surface of the Adsorba^®^ 300C cartridge. Surface-induced conformational changes in C3 promote alternative pathway amplification, leading to generation of C3a and C5a and deposition of C3b on the sorbent matrix. The anaphylatoxins trigger pulmonary intravascular macrophages, platelets, and neutrophils, resulting in thromboxane A_2_ release (measured as TXB_2_), pulmonary vasoconstriction, systemic hypotension, leukopenia, and hemoconcentration. The early C3a peak followed by delayed TXB_2_ elevation indicates sequential activation of humoral and cellular arms of innate immunity, consistent with a complement-driven mechanism of shock.

Overall, these findings extend the CARPA concept, as they suggest a cause-effect relationship between C activation and anaphylaxis. In this context, CARPA stands for C activation-related pseudoanaphylaxis.

### 3.6. Effects of oxygenator and dialysate cartridges on blood cells

Table 2 shows a decrease in all counted blood cell types immediately after initiation of extracorporeal circulation (i.e., after 0 min), most likely reflecting hemodilution due to expansion of the circulating blood volume. However, during the subsequent circulation period, none of the cell counts exhibited a consistent temporal trend.

**Table 2.** Blood cell changes caused by oxygenator and dialysate cartridges during extracorporeal circulation in pigs.

| Cartridge* | Minutes | WBC<br>10<sup>9</sup>/l | LYM<br>10<sup>9</sup>/l | NEU<br>10<sup>9</sup>/l | RBC<br>10<sup>12</sup>/l | HB<br>g/l | HCT<br>% | PLT<br>10<sup>9</sup>/l |
| --- | --- | --- | --- | --- | --- | --- | --- | --- |
| Oxygenator 1 | 0 | 10.7 | 8.3 | 2.2 | 4.5 | 68.0 | 24.2 | 328 |
|  | 15 | 7.6 | 7.0 | 0.6 | 3.7 | 55.0 | 19.9 | 204 |
|  | 30 | 8.3 | 7.1 | 1.1 | 3.5 | 54.0 | 18.7 | 196 |
|  | 45 | 8.6 | 7.3 | 1.2 | 3.5 | 55.0 | 18.9 | 223 |
|  | 60 | 10.3 | 8.5 | 1.8 | 3.9 | 59.0 | 20.8 | 282 |
| Oxygenator 2 | 0 | 17.67 | 14.48 | 3.01 | 5.31 | 85 | 28.5 | 320 |
|  | 15 | 11.44 | 10.65 | 0.72 | 4.25 | 69 | 22.9 | 211 |
|  | 30 | 13.18 | 11.32 | 1.67 | 4.42 | 68 | 23.5 | 225 |
|  | 45 | 13.09 | 11.47 | 1.5 | 3.97 | 64 | 21.4 | 200 |
|  | 60 | 14.17 | 12.15 | 1.91 | 4.33 | 67 | 23.3 | 207 |
| Dialysis filter | 0 | 13.7 | 8.6 | 4.9 | 5.0 | 82.0 | 25.8 | 327 |
|  | 15 | 10.8 | 7.3 | 3.4 | 4.4 | 69.0 | 23.0 | 300 |
|  | 30 | 8.5 | 6.8 | 1.7 | 4.4 | 71.0 | 22.8 | 321 |
|  | 45 | 7.6 | 6.4 | 1.2 | 4.5 | 70.0 | 23.7 | 309 |
|  | 60 | 11.9 | 7.2 | 4.6 | 4.5 | 74.0 | 24.1 | 322 |
| Oxygenator 3 | 0 | 17.7 | 12.8 | 4.8 | 5.6 | 91.0 | 30.0 | 254 |
|  | 15 | 14.2 | 11.3 | 2.9 | 4.8 | 77.0 | 20.0 | 190 |
|  | 30 | 14.4 | 11.4 | 2.9 | 4.8 | 77.0 | 20.0 | 202 |
|  | 45 | 14.3 | 11.5 | 2.7 | 4.7 | 77.0 | 20.0 | 216 |
|  | 60 | 15.3 | 12.0 | 3.3 | 4.8 | 77.0 | 30.0 | 203 |

\*The functions, composition and other specifics of different cartridges are given in Table 3 below.

## 4. Discussion

### 4.1. Toxicological concerns of extracorporeal cartridges

Table 3 compares the physicochemical characteristics and immune reactivity of the three cartridges used in the present study.

**Table 3.** Physicochemical parameters and immune reactivity of the extracorporeal cartridges.

| Parameter | Adsorba® 300C<br>adsorption column | MultiFiltratePRO<br>hemofilter | Inspire 8 Start P<br>oxygenator |
| --- | --- | --- | --- |
| Primary function | Adsorption of<br>toxins/cytokines | Diffusion/convection of<br>solutes | Gas exchange (O<sub>2</sub>/CO<sub>2</sub>) |
| Column matrix | cellulose-coated carbon<br>(charcoal) microparticles | Polysulfone/PES | Polypropylene |
| Surface area for<br>blood protein<br>binding | ~200,000-400,000 m<sup>2</sup>* | ~1.5–2.0 m<sup>2</sup> membrane | ~1.4–2.0 m<sup>2</sup> membrane |
| Geometric bed<br>contact surface area | 300 m<sup>2</sup> |  |  |
| Complement<br>activation | Very strong, rapid | Low–moderate | Strong, universal<br>during CPB |
| Subclinical<br>complement<br>activation | ~100% | Common | ~100% during CPB |
| Incidence of severe<br>anaphylactoid<br>reaction | <1% | ~0.02–0.05% | ≤0.1% |
| References | [20-24] | [25-27] | [28-30] |

\*This enormous surface area arises from micropores (< 2 nm) and mesopores (2-50 nm) inside the carbon particles which allow the passing of proteins such as C3/C3a)

As outlined in the Introduction, Adsorba^®^ 300C is a highly microporous, carbon-based adsorbent designed for extracorporeal blood purification to remove toxic or bioactive substances (e.g., in toxin or drug overdose). Due to safety concerns, it was withdrawn from commercial distribution as of 2022 [31]. The multiFiltratePRO polysulfone (or polyethersulfone, PES) hollow-fiber hemofilter also tested in this study is widely used for hemofiltration in acute kidney injury and represents standard-of-care therapy in ICU dialysis worldwide. Although substantially more biocompatible than earlier cellulose-based dialyzers, it can still cause “first-use” reactions in approximately <0.02–0.1% of dialysis sessions, primarily due to mild alternative pathway C activation [25-27]. The Inspire 8 Start P is a polypropylene hollow-fiber membrane oxygenator used in cardiopulmonary bypass (CPB). Contact of blood with the oxygenator surface has been shown to universally activate C, leukocytes, and other inflammatory cascades, contributing to postoperative systemic inflammation. Nevertheless, clinically overt hypersensitivity reactions during CPB are also very rare (0.01–0.1%) [28-30].

### 4.2. Complement activation as a cause of extracorporeal reactions

Exposure of blood to artificial surfaces, such as polymer-coated materials, usually cause C activation, an inherent feature of the innate immune system that evolved to recognize and eliminate “non-self” structures rapidly and without prior sensitization. Its broad sensitivity to unusual molecular patterns underlies the propensity of artificial materials to trigger this response [32-35].

One consequence of C activation is the release of anaphylatoxins (C3a, C5a), which induce allergy-like, so-called pseudoallergic reactions, whose most severe manifestation is (pseudo)anaphylaxis. Complement activation entails activation of blood cells (e.g., leukocytes, platelets) with liberation of numerous vasoactive and inflammatory mediators and enzymes that extend and amplify the initial anaphylatoxin signal. For this reason, the severity of pseudoallergic reactions is related to, but not necessarily proportional with the extent of C activation; hence the term complement activation-related pseudoallergy (CARPA). Although C activation is an inherent and nearly universal consequence of blood contact with foreign surfaces, the clinical manifestations of CARPA are unpredictable and typically rare, depending on multiple host-and device-related factors. The prediction of such rare reactions requires relevant animal models, however, modeling rare hypersensitivity reactions is difficult unless the animal species is intrinsically sensitive to the underlying mechanism.

### 4.3. The pig model of extracorporeal reactions

Pigs are uniquely suited to detect and quantify CARPA as they are naturally sensitive to C-induced anaphylactoid reactions due to the abundance of pulmonary intravascular macrophages and their well-characterized hemodynamic responsiveness to C-induced inflammatory mediators, most importantly thromboxane A2 [13-18]. Moreover, the body size of pigs permits the use of clinically approved human hemoperfusion cartridges, thereby enhancing the translational relevance of the findings.

With this rationale, in a previous study we used the pig CARPA model to compare the in vivo reactogenicity of three different dialysis cartridges [36]. We detected signs of mild CARPA, but only when blood from the extracorporeal circuit was reinfused into the animal [36]. It was hypothesized that reinfusion involved a brief stagnation of blood within the cartridge, allowing sufficient time for C3 to bind to the membrane surface and undergo cleavage to release C3a, the pivotal step in CARPA. These observations also implied that the total surface area of the cartridge membrane played a critical, rate-limiting role in these adverse reactions. Accordingly, a potentially useful strategy to prevent these reactions is the prevention of C activation on the membrane surface, rather than in the fluid phase, which may or may not be commensurate with the anaphylatoxin attack. Obviously, to engineer such a column, assessing CARPA as a function of column surface area is essential. In this regard, the pig model provided a unique opportunity, particularly given the availability of hemoperfusion columns with enormous surface area accessible for C3 binding.

### 4.4. The goal and message of the present study

To test the hypothesis that membrane surface area is a rate-limiting factor in CARPA, we extended our previous study [36] by using Adsorba^®^ 300C, a hemoperfusion column exposing an extremely high, 5 orders of magnitude greater surface area for interaction with C3, than that of the comparator dialysis and oxygenator cartridges (Table 3). The finding that Adsorba^®^ 300C, but not the dialyzer or oxygenator columns, induced an unprecedentedly severe and sustained anaphylactic shock, supports the surface area dependence of CARPA hypothesis. Accordingly, extracorporeal circulation with Adsorba^®^ 300C in pigs provides a valid animal model for studying rare but potentially severe, and occasionally lethal, extracorporeal circuit, as well as it provides a useful tool to develop C-neutralized columns. Furthermore, the temporal delay between the rise in C3a and the subsequent TXB_2_ response represents strong evidence for a causal relationship between C activation and TXA_2_ release, thereby strengthening the CARPA concept.

### 4.5. Clinical relevance of studying severe extracorporeal reactions

Extracorporeal reactions are generally considered rare; however, their absolute burden becomes clinically relevant when viewed against the enormous global exposure to extracorporeal artificial surfaces. Approximately 3-4 million patients worldwide receive maintenance hemodialysis, typically three times weekly, resulting in an estimated 550-650 million hemodialysis sessions annually [37-39]. Together with an estimated 1.5–2.2 million CPB procedures each year [40, 41], even assuming the lowest estimated incidence of severe anaphylaxis (e.g., 0.02%), these exposure numbers would translate into approximately 110,000–130,000 affected patients annually worldwide. With a reported mortality rate of ∼0.5-2% among hospitalized anaphylaxis cases [42, 43], this could correspond to around a thousand deaths a year. These figures underscore the substantial clinical relevance of developing effective preventive strategies against extracorporeal circuit-induced anaphylactic reactions, for which the pig CARPA model may provide a valuable translational platform.

### 4.6. A new large animal model of fatal anaphylactic shock

Beyond the clinical relevance of our findings in extracorporeal procedures, as outlined above, the observation of uniquely severe and sustained anaphylaxis also establishes a novel large-animal model of irreversible, fatal anaphylactic shock. Existing models using mice [44, 45], rats [46-48], and guinea pigs [49-51] predominantly reproduce severe but reversible cardiovascular and pulmonary reactions that resolve after withdrawal of the trigger or pharmacological intervention. In contrast, a defining feature of the present extracorporeal perfusion model in pigs is that the adsorbent cartridge provides continuous immune activation, thereby maintaining cardiovascular instability despite administration of standard rescue therapies. The repeated pattern of transient blood pressure recovery followed by rapid relapse into shock indicates failure of compensatory vascular and cardiac mechanisms and clearly distinguishes this condition from reversible CARPA, classical IgE-mediated anaphylaxis, or acute infusion reactions.

Across species, there is currently no standardized, clinically relevant model that consistently mimics sustained, self-perpetuating shock progressing to death despite resuscitative measures, representing a major gap in pharmacology and toxicology research. Thus, our model offers a novel approach to studying hemodynamic collapse driven by persistent anaphylatoxin generation, inflammatory mediator release, and blood cell activation and redistribution. Such immune-vascular collapse extends well beyond the context of extracorporeal reactions, thus the porcine model in this study provides a unique experimental platform for advancing the understanding, prevention, and treatment of a commonly occurring lethal acute condition.

### 4.7. The limitations of this study

Although the present study involved a single pig exposed to the Adsorba^®^ cartridge, the constellation, timing, and severity of the observed hemodynamic and hematologic changes closely mirrored both the previously reported CARPA caused by reactive nanoparticles [13, 14, 16-18], and the repeatedly documented adverse reactions in humans undergoing hemoperfusion (Table 1) [3-10] Such a complete phenotypic overlap makes random variation, experimental artifact, or an outlier animal an unlikely explanation for the findings. In particular, the exceptionally severe and prolonged anaphylactic reaction observed in only 1 of 5 pigs, specifically in the animal exposed to the highly immune-reactogenic cartridge, provides proof of concept for the applicability of this model to the study of human extracorporeal reactions. The inclusion of additional animals would allow a more precise characterization of inter-individual variability and cartridge dependency of these reactions, however, the primary objective of this pilot study was not statistical generalization but proof of principle, namely to demonstrate that the porcine CARPA model may serve as a platform technology for testing therapeutic and prophylactic interventions of extracorporeal anaphylaxis.

## 5. CRediT authorship contribution statement

Conceptualization, JSz, BAB, TR, JAJ; methodology and formal analysis SS, AJH, ABD, LM, TM, GTK, RF LD, JN; writing/editing: JSz, BAB, TR, JAJ, LM; project administration and funding, TJ, BM, JAJ, All authors have read and agreed to the published version of the manuscript.

## 7. Review Board Statement

The study was conducted in accordance with the Declaration of Helsinki, and the animal study protocol was approved by the Institutional and National Review Board/Ethics Committees, PE/EA/843-7/2020.

## 8. Data Availability Statement

Experimental data are available upon reasonable request to the corresponding author.

## 9. Acknowledgments

The expert technical support by Dóra Szkrajcsics, Maria H. Velkei, Katalin Simay, Henriett Biró and Krisztina Fazekas are gratefully acknowledged. ORCID (JS): 0000-0003-1738-797X, (JAJ): 0000-0002-1800-8102 Conflict of interest: None of the authors declared conflict of interest

## 10. Declaration of transparency and scientific rigor

The authors acknowledge that this paper adheres to the principles for transparent reporting and scientific rigour of preclinical research as stated in the BJP guidelines for Natural Products Research, Design and Analysis, Immunoblotting and Immunochemistry, and Animal Experimentation, and as recommended by funding agencies, publishers and other organizations engaged with supporting research.

